# Elevated Hoxb5b expands vagal neural crest pool and blocks enteric neuronal development in zebrafish

**DOI:** 10.1101/2021.10.28.466356

**Authors:** Aubrey G. Adam Howard, Aaron C. Nguyen, Joshua Tworig, Priya Ravisankar, Eileen W. Singleton, Can Li, Grayson Kotzur, Joshua S. Waxman, Rosa. A. Uribe

## Abstract

Neural crest cells (NCCs) are a migratory, transient, and multipotent stem cell population essential to vertebrate embryonic development, contributing to numerous cell lineages in the adult organism. While great strides have been made in elucidating molecular and cellular events that drive NCC specification, comprehensive knowledge of the genetic factors that orchestrate NCC developmental programs is still far from complete. We discovered that elevated Hoxb5b levels promoted an expansion of zebrafish NCCs, which persisted throughout multiple stages of development. Correspondingly, elevated Hoxb5b also specifically expanded expression domains of the vagal NCC markers *foxd3* and *phox2bb*. Increases in NCCs were most apparent after pulsed ectopic Hoxb5b expression at early developmental stages, rather than later during differentiation stages, as determined using a novel transgenic zebrafish line. The increase in vagal NCCs early in development led to supernumerary Phox2b^+^ enteric neural progenitors, while leaving many other NCC-derived tissues without an overt phenotype. Surprisingly, these NCC-derived enteric progenitors failed to expand properly into sufficient quantities of enterically fated neurons and stalled in the gut tissue. These results suggest that while Hoxb5b participates in vagal NCC development as a driver of progenitor expansion, the supernumerary, ectopically localized NCC fail to initiate expansion programs in timely fashion in the gut. All together, these data point to a model in which Hoxb5b regulates NCCs both in a tissue specific and temporally restricted manner.

## Introduction

As an embryonic stem cell population in vertebrates, neural crest cells (NCCs) are renowned for their remarkable migratory capacity, as well as their multipotency. Born from the dorsal neural tube, NCCs migrate along stereotypic routes throughout the early embryo and give rise to a wide range of diverse tissue lineages, such as craniofacial skeleton, portions of the peripheral nervous system, and pigment cells (Rocha et al., 2020). NCCs exhibit regional potential along the anteroposterior (AP) neuraxis such that they may be divided into four general populations: cranial, vagal, trunk, and sacral (Le Douarin, 1982; Le Douarin and Kalcheim, 1999). Each of these populations give rise to numerous discrete lineages, for example, cranial NCC largely give rise to cell lineages in the head. Particularly of interest are vagal NCCs, which contribute to several tissues, such as the cardiac outflow tract and nearly all of the enteric nervous system (ENS) (Tang et al., 2021) within the gut, and have been less well characterized than other populations (Hutchins et al., 2018). While the driving genetic factors which regulate the general pattern of NCC developmental trajectories have been well described (Martik and Bronner, 2017; Simoes-Costa and Bronner, 2016), we still have an incomplete understanding of what genes function in context of vagal NCC development and their subsequent differentiation.

One factor that has been implicated in vagal NCC development is Hoxb5. In mice, dominant negative abrogation of embryonic Hoxb5 activity alters vagal and trunk NCC development, with reduced NCCs observed en route to and along gut tissue, as well as decreased numbers of melanoblasts throughout the body (Kam and Lui, 2015; Lui et al., 2008). Additionally, Hoxb5 may regulate expression of key genes active in NCC development, notably *Foxd3* (Kam et al., 2014a) and *Phox2b* (Kam and Lui, 2015). The orthologous gene in zebrafish, *hoxb5b*, which is the primary ortholog in teleost fishes (Jarinova et al., 2008), was detected in differentiating NCC lineages at both 48 and 68 hours post fertilization (hpf) (Howard et al., 2021), suggesting it may also play a role during zebrafish NCC development. While characterization of zebrafish *hoxb5b* mRNA expression *in situ* has been pervasively characterized in several early embryonic contexts, such as the mesoderm and limb buds (Hortopan and Baraban, 2011; Jarinova et al., 2008; Kudoh et al., 2001; van der Velden et al., 2012; Waxman et al., 2008), the functional role of *hoxb5b* with respect to NCC development had not yet been examined.

Here, we postulated that *hoxb5b* functions as a potential driver of vagal NCC development. We provide evidence that overexpression of *hoxb5b* is sufficient to grossly expand NCC populations throughout the embryo, in addition to ectopic expansion of vagal domains marked by *foxd3* and *phox2bb*. The functional window of Hoxb5b activity was also restricted to a narrow developmental span, early during embryogenesis, rather than during NCC differentiation stages. The early expansion of NCC, however, did not lead to corresponding pan increases in NCC-derived tissues. Rather, elevated Hoxb5b activity expanded enteric neural progenitor cell pools along the gut, yet suppressed their subsequent expansion as they differentiated into enteric neurons, leading to overall fewer neurons along the gut. These data cumulatively support a model in which *hoxb5b* is a potent regulator of NCC expansion and cell number in zebrafish.

## Results

### *hoxb5b* is expressed in post-otic vagal NCCs during zebrafish development

We examined *hoxb5b* expression in zebrafish embryos (**Figure 1A**) using whole mount *in situ* hybridization (ISH). At 32 hpf, *hoxb5b* was expressed bilaterally immediately posterior to the otic vesicle (post-otic), along the foregut, in the hindbrain, and anterior spinal cord (**Figure 1B**), as previously described (Jarinova et al., 2008). *hoxb5b* persisted in all three domains at 50 hpf (**Figure 1C**), though the post-otic domains (POD) were slightly restricted and the hindbrain/spinal cord expression gained a more defined posterior boundary. By 77 hpf, *hoxb5b* expression remained largely in the hindbrain and foregut, with diminished yet persistent expression in the POD (**Figure 1D**). Together, these ISH data reveal changing post-otic spatiotemporal expression patterns of *hoxb5b* during the first four days of development.

**Figure 1.**
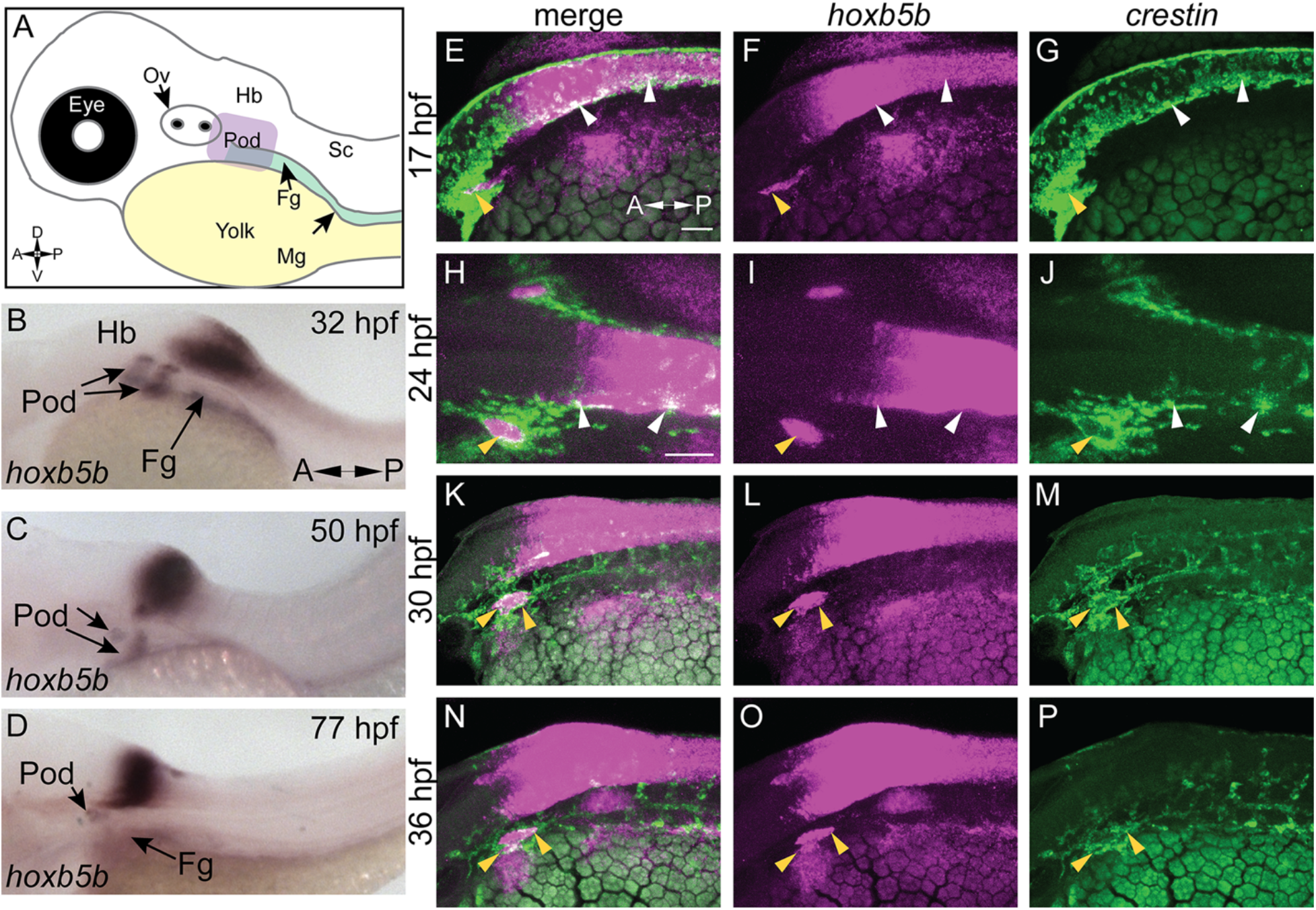
hoxb5b is expressed during early NCC development. **(A)** Schematized embryo illustrating approximate locations of major relevant anatomical features, namely the post-otic domain (POD), eye, yolk, Otic Vesicle (Ov), the Hindbrain (Hb), presumptive spinal cord (Sc), foregut mesenchyme (Fg), and Midgut mesenchyme (Mg). **(B-D)** In Situ Hybridization demonstrating *hoxb5b* expression in the posterior Hindbrain, Pod, and Fg during the second through third day of development. **(E-P)** Hybridization Chain Reaction probes against *crestin* and *hoxb5b* highlight their overlapping domains in the Pod (yellow arrowheads) and along the dorsal length (white arrowheads). Scale bars E,H: 50 μM.

We next examined the relationship of *hoxb5b* expression to the vagal NCC population during NCC specification and migration phases. NCCs were assayed by using the zebrafish pan-NCC marker *crestin* (Luo et al., 2001), in combination with *hoxb5b*, via hybridization chain reaction (HCR) (Choi et al., 2010, 2016, 2018). At 17 hpf, *crestin*^*+*^/*hoxb5b*^+^ cells were present dorsally (**Figure 1E-G, white arrowheads**), as well as ventral-laterally, along a post-otic stripe (**Figure 1E-G, yellow arrowhead**), revealing that *hoxb5b* is expressed within the POD vagal NCC population. *crestin*^*+*^/*hoxb5b*^+^ regions persisted by 24 hpf dorsally, in posterior hindbrain/anterior spinal cord axial levels (**Figure 1H-J, white arrowheads**). Concurrently at this stage, *hoxb5b* expression within the stripe became internalized within the POD vagal NCC population, marking the central cells of this region and highlighting several NCCs which expressed *hoxb5b* (**Figure 1H-J, yellow arrowhead**). Between 30-36 hpf, the stripe co-positive *hoxb5b*^*+*^/*crestin*^*+*^ stripe of POD vagal NCC persisted (**Figure 1K-P, yellow arrowheads**), and *crestin*^*+*^/*hoxb5b*^+^ cells were still observed along the dorsal neural tube. These data are consistent with our prior findings in which *hoxb5b* mRNA was present in posterior neural crest at 48 hpf and 68-70 hpf (Howard et al., 2021). Collectively, the ISH and HCR data indicate that *hoxb5b* mRNA expression is coincident with a subset of vagal NCCs, persisting throughout the developmental window spanning NCC specification and well into their migration phase, highlighting *hoxb5b*’s potential role as a driver of vagal NCC development.

### Elevated Hoxb5b activity globally promotes expanded localization and increased number of NCC *in vivo*

While other work has focused exclusively on the loss of function of Hoxb5 related genes (Dalgin and Prince, 2021; Lui et al., 2008), we have sought to understand the gain of function role of *hoxb5b* in the context of NCC development. To examine the possible role of *hoxb5b* in NCC, we employed a hyperactive *vp16-hoxb5b* fusion construct (Waxman and Yelon, 2009; Waxman et al., 2008). Injection of *vp16-hoxb5b* mRNA resulted in expansion of NCC, when compared to control embryos at 32 hpf, as assayed with an ISH probe against *crestin* (**Figure 2A**,**B**). The expansion in NCC territory was prominent in the pre-otic region (**Figure 2B; white arrowheads**), the POD (**Figure 2B; yellow arrowheads**) and along the spinal cord-level of the trunk (**Figure 2B; red arrowheads**). Strikingly, post-otic expansion persisted along the dorsal-ventral and anterior-posterior axes well through NCC migration phases at 50 hpf (**Figure 2C**,**D; arrowheads**). Furthermore, quantification of the area occupied by POD NCCs corroborated the observed expansion of NCC localization (**Figure 2E**).

**Figure 2.**
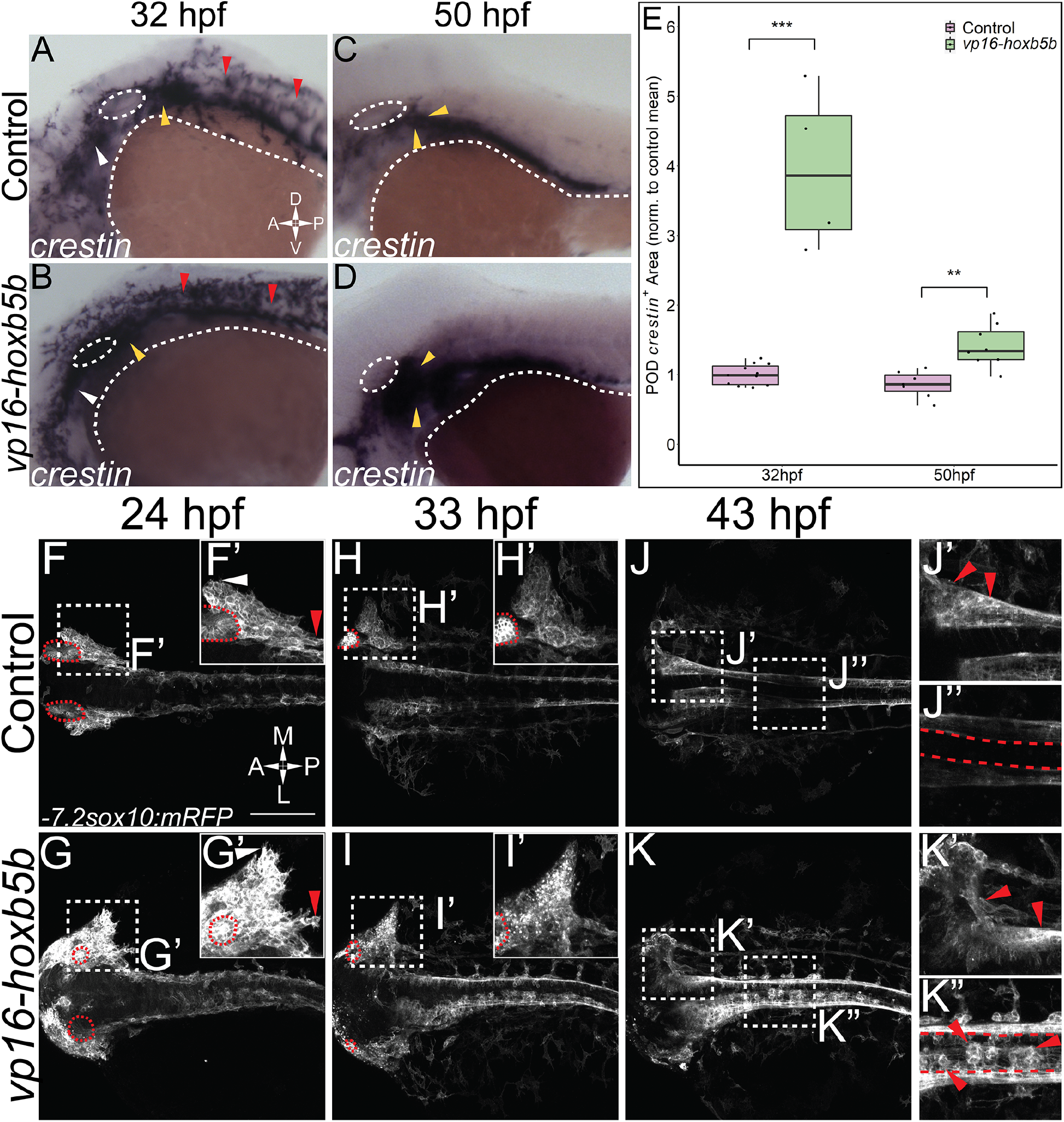
Elevated Hoxb5b activity globally increases both number and localization of neural crest cells. **(A-D)** In Situ Hybridization for NCC using a *crestin* probe at both 32 hpf (A,B) and 50 hpf (C,D). *crestin*^*+*^ domains for embryos injected with 15 pg of *vp16-hoxb5b* mRNA (B,D) were expanded in the post-otic (yellow arrowheads), cranial (white arrowheads), and spinal cord (red arrowheads) compared to uninjected embryos (A,C). **(E)** Quantification of expanded vagal *crestin*^+^ domains shows significant expansion at both 32 hpf (*p* = 0.75×10^−7^) and at 50 hpf (*p* = 0.00114). **(F-K)** Maximum intensity projected stills taken from a confocal acquired time lapse of movies of *sox10:mRFP* embryos. Controls were compared to 30 pg *vp16-hoxb5b* injected embryos, examined from 24 hpf to 43 hpf and serially imaged along the dorsal aspect of the vagal domain. mRFP^+^ NCCs are grossly expanded in the vagal domain (G,G’ arrowheads) over controls (F,F’, arrowheads). This expansion persists through the course of development, resulting in ectopically localized cells along the dorsal aspect of the embryo (K’’) and in the post-otic pool (K’). Scale Bars in F: 100μM. Anterior: Left

To better understand the spatiotemporal distribution of the increased NCC following *vp16-hoxb5b* expression, we utilized confocal microscopy to image *-7*.*2sox10:mRFP* transgenic embryos (referred to here as *sox10:mRFP*) (Kucenas et al., 2008), where NCC are labeled using a membrane bound RFP. Congruent with our prior findings (**Figure 2A, B**), at 24 and 33 hpf, confocal projections revealed *vp16-hoxb5b* expressing embryos exhibited broadened POD vagal NCC domains along the anterior-posterior and medio-lateral axes (**Figure 2G, G’, I, I’; arrowheads**), when compared to control embryos (**Figure 2F, F’, H, H’; arrowheads**). This expansion was coupled with an increase in the number of POD NCC. By 43 hpf, *vp16-hoxb5b* expressing embryos displayed a striking expansion of the POD NCC, as well as a disruption of the overall architecture of the domain, which extended further laterally from the dorsal midline (**Figure 2K’**), than in control (**Figure 2J’**). Moreover, *vp16-hoxb5b* promoted ectopic accumulation of cells along the dorsal midline of the spinal cord (**Figure 2K’’**), which were not observed in control embryos (**Figure 2J’**’). Considered together, these data (**Figure 2**) indicate that elevated Hoxb5b activity alters NCC patterning along the embryo as well as expanding their cell number.

### Elevated Hoxb5b activity is sufficient to expand vagal NCC marker expression

That *vp16-hoxb5b* expressing embryos contained an overabundance of NCC, and that the overproduced NCC were prominently enriched in the vagal axial levels suggests that excess Hoxb5b activity influences vagal NCC development in zebrafish. To examine if vagal NCC specification was altered following excess Hoxb5b activity, we assayed the expression of canonical marker genes of the vagal NCC, *foxd3* (Lister et al., 2006) and *phox2bb* (Elworthy et al., 2005). At 32 hpf, *foxd3* expression serves as an indicator of multipotent NCC while *phox2bb* indicates NCCs which are now specified to an autonomic neural lineage, particularly the ENS (Pattyn et al., 1999; Uribe and Bronner, 2015). *vp16-hoxb5b* expression was sufficient to widen *foxd3* expression domains at 32 hpf, principally along the anterior-posterior and mediolateral directions in the vagal region, when compared to control expression patterns (**Figure 3A**,**B; black bars**). We found that *vp16-hoxb5b* expression expanded the POD *foxd3*^+^ area, leading to a 1.95 fold increase in the mean domain size compared to controls (**Figure 3E**). Additionally, *phox2bb* was also greatly expanded in response to increased Hoxb5b activity along the hindbrain (**Figure 3C**,**D; black bars**), with expansion uniformly in both the anterior-posterior and mediolateral axis, similar to the expansion of the *foxd3*. Measuring the hindbrain *phox2bb*^*+*^ domain, we observed a 43% increase in the *phox2bb* expressed area throughout the hindbrain following elevated Hoxb5b (**Figure 3F**). Lastly, POD *phox2bb* expression was dramatically altered by elevated Hoxb5b (**Figure 3C**,**D; white arrow heads**), with a 2.14 times mean increase in domain size compared to wild-type controls (**Figure 3G**). In all, these data indicate that elevated Hoxb5b activity drastically expands vagal NCC marker expression along the embryo.

**Figure 3.**
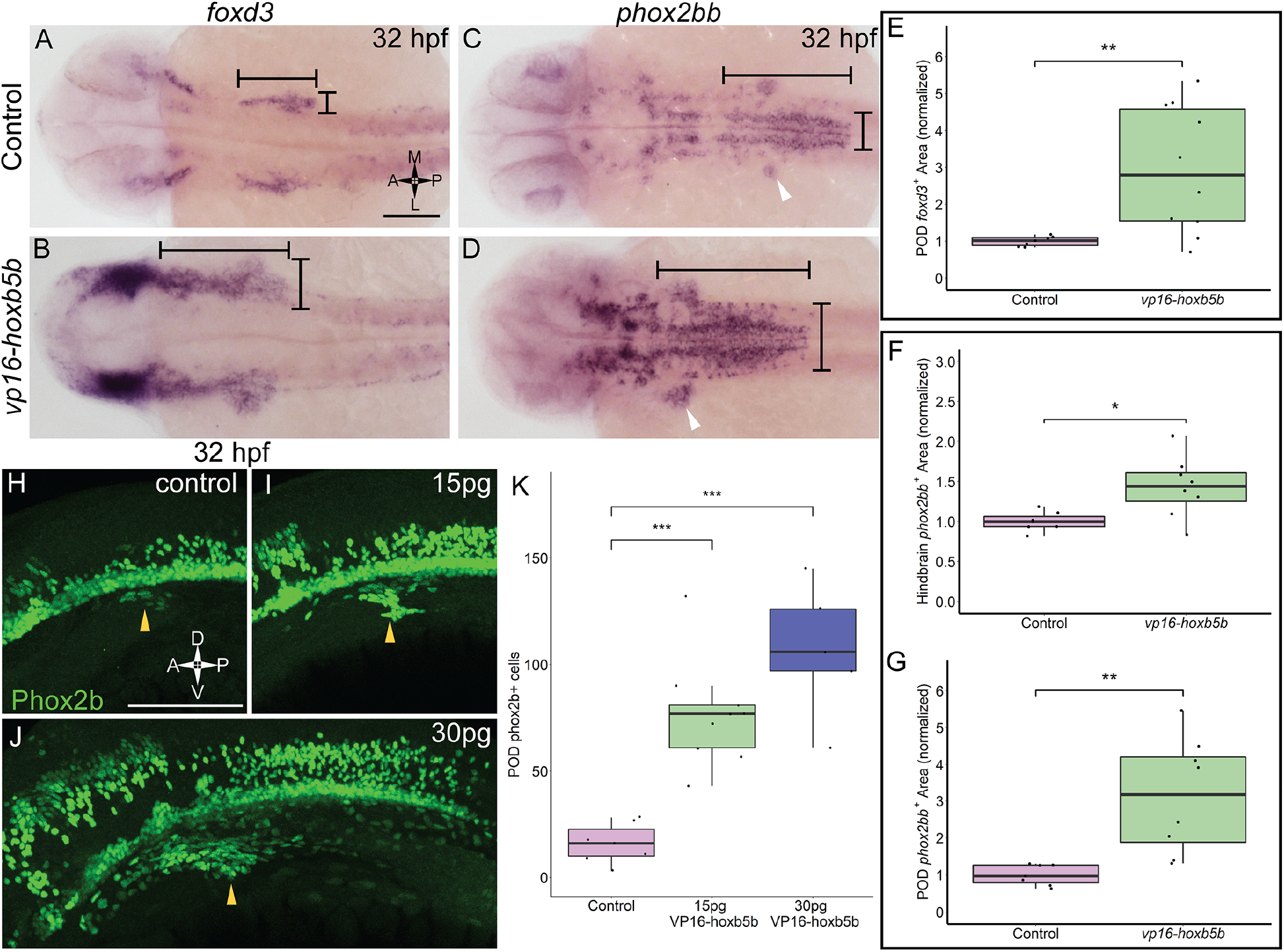
Elevated Hoxb5b activity expands the expression domains of vagal markers *foxd3* and *phox2bb* during the first day in development. **(A-B)** ISH against *foxd3*, a marker for multipotent NCCs at this stage, in 32 hpf. Embryos injected with 30 pg *vp16-hoxb5b* were compared to WT controls. **(C-D)** ISH against *phox2bb*, an autonomic NCC marker, in 32 hpf. Embryos injected with 30 pg *vp16-hoxb5b* were compared to WT controls. The POD is denoted with a white arrow head. **(E-G)** Quantified POD *foxd3* expression (E, *p* = 0.00571) or *phox2bb* expression (F, *p* = 0.138), as noted in representative images by black bars. The discrete post otic expression domain is quantified in (G, *p* = 0.00563). Areas are normalized to control mean. **(H-K)** IHC detection of Phox2b^+^ cells viewed along the lateral axis of 32 hpf embryos reveal that WT controls already have a nascent population (H, white arrowheads). *vp16-hoxb5b* overexpression, at either 15 pg (I) or 30 pg (J) of mRNA injected, expands the Phox2b^+^ vagal NCCs (white arrowheads). (K) Counted Phoxb^+^ cells from 3 dimensional micrographs reveal increasing cells with the amount of *vp16-hoxb5b* mRNA (15 pg: p = 3.03 × 10^−5^; 30 pg: p = 2.61 × 10^−5^). Scale bar A,L: 100 μM

In support of the specific expansion of POD localized *phox2bb* expression, we also observed a corresponding increase in the number of Phox2b^+^ cells, via whole mount fluorescent Immunohistochemistry (IHC), using an antibody against Phox2b (**Figure 3-Supplement 1A-C; Figure 3)**. At 32 hpf, Phox2b^+^ cells were observed in the POD, with 16 cells on average (**Figure 3L; arrowhead, Figure 3K**). Overexpression of Hoxb5b stimulated a dramatic expansion of POD Phox2b^+^ cells (**Figure 3I**,**J; arrowheads**). The increase in POD Phox2b^+^ cells was also concordant with an increase in *vp16-hoxb5b* dosage, with 77 and 107 mean POD Phox2b^+^ cells per animal detected following injection with either 15 pg or 30 pg of mRNA, respectively (**Figure 3K**). The quantifiable increase in POD Phox2b^+^ cells is confirmatory of our prior qualitative observations regarding increased cell number (**Figure 2**) and positions Hoxb5b as a potent driver of NCC number and localization.

### Hoxb5b overexpression increases NCC production during early NCC development

While we observed supernumerary NCCs ectopically localized, and expanded expression of vagal NCC specification factors following global expression of *vp16-hoxb5b* mRNA, exactly when during NCC development Hoxb5b may exert its influence was still unclear. To investigate the potential temporal role(s) of Hoxb5b during NCC development, we created and utilized a novel transgenic line, Tg(*hsp70l:EGFP-hoxb5b;cryaa:dsRed*)^ci1014^ (hereafter referred to as *hsp70l:GFP-hoxb5b*), which enables pan ectopic expression of a GFP-Hoxb5b protein fusion under the thermally inducible *hsp70l* promoter (Kwan et al., 2007) (**Figure 4A**). The transgene can then be activated by rapidly transferring embryos to warm 37°C culture conditions, which drives strong global expression of the EGFP-Hoxb5b fusion protein throughout the embryonic tissues (**Figure 4 – Supplement 1A**). EGFP-Hoxb5b demonstrated robust and distinctive nuclear localization, which was still detectable over 24 hours after embryos were returned to 28°C (**Figure 4-Supplement B-C**).

**Figure 4.**
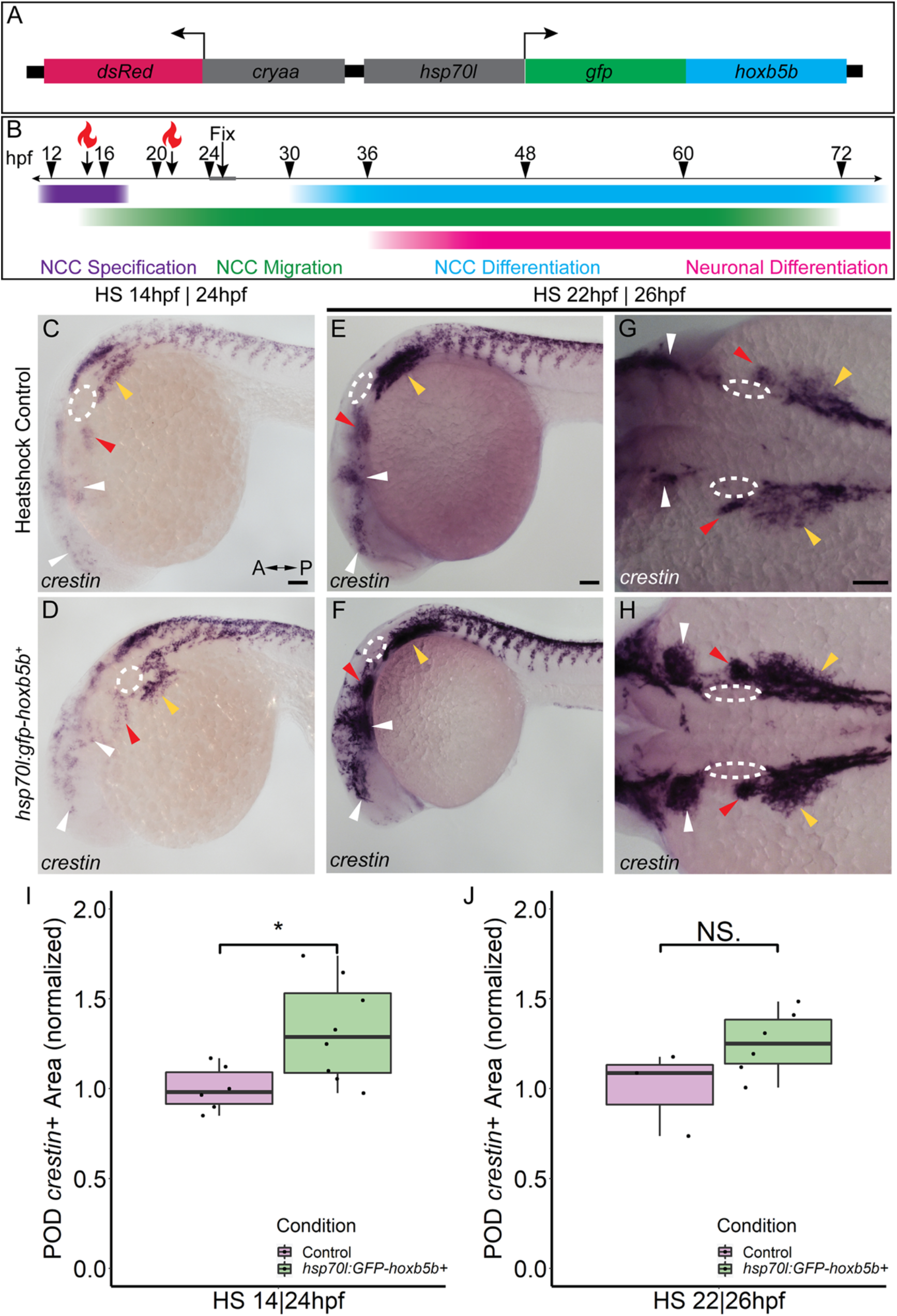
Temporally controlled overexpression of Hoxb5b during the first day in development are sufficient to expand the neural crest pool. **(A)** Schematized model of the *hsp70l:gfp-hoxb5b;cryaa:dsRed* genetic construct. **(B)** Illustration depicting specific periods of heat shock for embryo groups relative to classical hallmarks of NCC development. **(C-D)** ISH using a probe for *crestin* in *hsp70l:GFP-hoxb5b*^+^ embryos heat shocked at 14 hpf and fixed at 24 hpf, compared to GFP^-^ sibling controls treated in parallel. Dramatic expansion of the POD can be seen (yellow arrowheads) in the Hoxb5b overexpressing embryos, while more subtle expansion is noted in the cranial NCC (white arrowheads) and pre-otic crest (red arrowheads). **(E-H)** Similar to (C-D) ISH with a *crestin* probe in *hsp70l:gfp-hoxb5b* and GFP-sibling controls heat shocked at 22 hpf and fixed for ISH at 26 hpf, examining both *crestin*^*+*^ domains both laterally (E-F) and dorsally (G-H). **(I-J)** Graphs depicting areas occupied by *crestin* staining in both GFP-control and Hoxb5b overexpressing embryos. Hoxb5b overexpression at 14 hpf was sufficient to expand NCC localization and qualitatively NCC number (I, *p* = 0.0234). A later heat shock at 22 hpf – fixation at 26 hpf, did not expand vagal NCC localization (J, *p =* 0.1125), but did appear to increase *crestin* staining, indicative of an increase in NCC number. Scale bars C,E,G: 100 μM

Two early phases in NCC development were tested, with heat shocks conducted starting either at 14 hpf, during NCC specification, or 22 hpf, early during the migratory span of posterior NCCs (**Figure 4B**). After heat shock at 14 hpf, whole mount ISH at 24 hpf showed enlargement of *crestin*^*+*^ NCC domains in GFP-Hoxb5b^+^ embryos, over the GFP-Hoxb5b^-^ sibling controls (**Figure 4C-D**), particularly prevalent in the POD NCC (**Figure 4D; yellow arrowheads**). This increase in area was also accompanied by a qualitative increase in the number of *crestin*^*+*^ cells along the posterior dorsal length of the embryo, similar to the phenotype observed in the previous Hoxb5b mRNA overexpression assays (**Figure 2A-E**). Quantified area of *crestin*^+^ PODs confirmed the expansion of vagal NCCs (**Figure 4I**). Additionally, subtle expansion of cranial NCCs (**Figure 4C-D, white arrowheads**) and pre-otic NCCs (**Figure 4C-D, red arrowheads**) was also observed, though far less striking than that of the vagal population at this stage. These data indicate that NCC localization during early specification phase of NCC development is receptive to Hoxb5b activity.

GFP-Hoxb5b induction at 22 hpf also increased *crestin* staining by 26 hpf throughout the POD (**Figure 4E-H, yellow arrowheads**), which is especially prominent in the NCCs most proximal to the otic vesicles (**Figure G-H, red arrowheads**). As in the heat shock at 14 hpf, increased NCC localization was also observed in GFP-Hoxb5b^+^ embryos across the cranial NCC populations (**Figure 4E-H; white arrowheads**). Interestingly, while this later heat shock did not significantly expand measurable vagal NCC area (**Figure 4J**), it did increase their abundance (**Figure E-H**). Nonetheless, this subtle difference in only a few hours of development indicates a critical period during early NCC development in which Hoxb5b is sufficient to promote NCC localization in the developing vertebrate body.

### Increased Hoxb5b alters specific vagal NCC-derived tissues

Due to the multipotent nature of NCC, it is possible a wide diversity of tissues can be affected by even a small perturbation in NCC development. Because we observed an overproduction of NCC following increases in Hoxb5b activity, we wondered if downstream NCC derivatives were also affected. Therefore, we first investigated the effect of *vp16-hoxb5b* expression on vagal and trunk NCC-derived cell types; including, pigment cells (melanophores and iridophores) and neuronal derivatives. Embryos expressing *vp16*-*hoxb5b* were able to produce pigment cells without appreciable differences (**Figure 5 – Supplement 1 A-D**). Both dorsal root ganglia (DRG) and Superior Cervical Ganglion (SCG) are derived from the vagal/trunk NCC pool (Durbec et al., 1996). Neurons comprising the DRG and SCG were largely normal following *vp16-hoxb5b* injection (**Figure 5 – Supplement 1 E-H**).

We next assayed the NCC-derived cell population along the gut length (**Figure 5A**), enteric neural progenitors, which typically migrate to the midgut level by 2 dpf and will have colonized the hindgut by 3 dpf, giving rise to neurons of the enteric nervous system (Ganz, 2018). We found that by 50 hpf, Phox2b^*+*^ enteric neural progenitors significantly increased after *vp16*-*hoxb5b* expression (**Figure 5C-D**), compared to the controls (**Figure 5B**). Counting Phox2b^+^ cells in the POD and along the gut tract (**Figure 5B-D; white dashes**), revealed the number of cells trended with increasing amount of *vp16*-*hoxb5b* mRNA injected (**Figure 5E**), consistent with the phenotype at 32 hpf (**Figure 3H-J**). The supernumerary enteric neural progenitors were observed together with their accumulation along the foregut (**Figure 5B-D; yellow arrowheads**) and in the POD (**Figure 5F**), though an increase in the number of cells was also observed at the end of the enteric migration chain along the midgut (**Figure 5 – Supplement 1I; Figure 5C**,**D; white arrowheads**). The increase in cells was uniform across the gut tract, with no change in the fraction of Phox2b^+^ cells found in the POD or gut mesenchyme after Hoxb5b perturbation (**Figure 5 – Supplement 1J**). These findings indicate that elevated Hoxb5b elicits a global increase in enteric neural progenitor number through the first two days of development.

**Figure 5.**
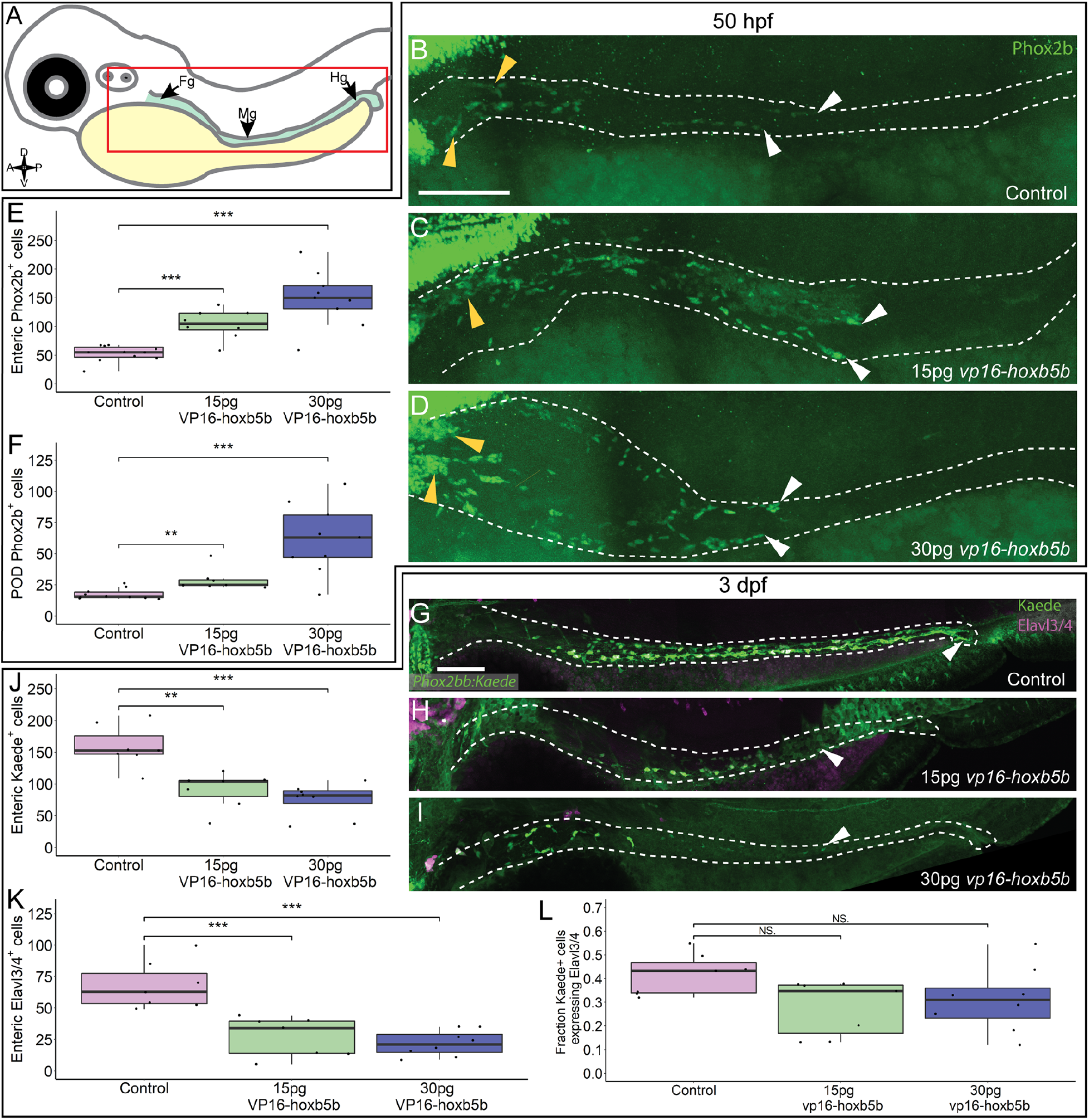
Hoxb5b is sufficient to expand early enteric neural progenitors. **(A)** Schematized model of a zebrafish embryo highlighting the region of the gut tube, which is imaged in the following panels. **(B-D)** Whole mount IHC for Phox2b in 50 hpf control embryos (B) compared to embryos injected either with 15 pg (C) or 30 pg (D) of *vp16-hoxb5b* mRNA. Yellow arrow heads indicated POD localized Phox2b^+^ cells, white arrows designate terminal end of enteric NCC chain, which falls within the gut tract outlined with white dashes. **(E-F)** Quantified cell numbers reveal a coordinate increase in Phox2b^+^ along the gut axis at 50 hpf trending with increasing *vp16-hoxb5b* mRNA amounts (E, 15 pg: *p* = 2.834 × 10^−5^; 30 pg: *p* = 0.00318). Additionally, the number of Phox2b^+^ cells restricted to the POD also increased in response to elevated Hoxb5b activity (F, 15 pg: *p* = 0.00328; 30 pg: *p* = 0.00068). **(G-I)** Whole mount IHC on -8.3*phox2bb:kaede* embryos with antibodies against Elavl3/4 and Kaede, marking the enteric NCC lineage cells. **(J-L)** Quantification of the number of enteric neural progenitors (J, 15 pg: *p* = 0.00011; 30 pg: *p* = 4.09 × 10^−5^) and differentiating enteric neurons (K, 15 pg: *p* = 0.001352; 30 pg: *p* = 0.0001042) at 3 dpf show decreasing numbers of both cell populations. However, the total fraction of differentiating (Hu^+^) NCC-derived Kaede^+^ cells unchanged following elevated Hoxb5b activity (L, 15 pg: *p* = 0. 0.1282; 30 pg: *p* = 0.102). Scale Bar B,G: 100 μM

To determine if the supernumerary enteric cells were capable of differentiating into neurons later in development, we utilized *-8*.*3phox2bb*:*kaede* transgenic embryos which label enteric progenitors during their early neuronal differentiation (Harrison et al., 2014). Surprisingly, despite the increase in enteric progenitors at 50 hpf, enteric neurons by the 3 dpf were dramatically decreased in *vp16-hoxb5b* expressing embryos compared to controls. Kaede^+^ cells successfully colonized the gut length by 3 dpf in control embryos (**Figure 5G; white arrowhead**), many cells of which (42%) also co-expressed the pan neuronal marker Elavl3/4 (**Figure 5G**,**K**,**L**), signaling the onset of neuron differentiation. In contrast, *vp16-hoxb5b* expressing embryos at both doses displayed a drastic loss of Kaede^+^ and Elavl3/4^+^ enteric cells (**Figure 5J**,**K**), with the remaining Kaede^+^ cells failing to localize past the level of the midgut (**Figure 5H**,**I; white arrowheads**). The fraction of Elavl3/4^+^/Kaede^+^ cells in both *vp16-hoxb5b* expressing conditions were reduced, at .31 and .28 respectively, when compared to control at .42 (Figure 5L). While not reaching significance, when these data are taken together with the significant reduction in total enteric cell numbers along the gut (**Figure 5J**,**K**), they likely indicate that general enteric progenitor pool depletion along the gut affects subsequent proper numbers of enteric neurons, following elevated Hoxb5b expression. Overall, these results suggest that while supernumerary enteric neural progenitors are present at and before 2 dpf after elevated Hoxb5b, they largely depleted by the 3 dpf.

### Hoxb5b influences enteric patterning during early developmental stages

In order to ascertain the timing during which excess Hoxb5b activity affects enteric nervous system development, we again leveraged the *hsp70l:GFP-hoxb5b* fish line. Embryos were heat shocked during NCC specification (14 hpf), migration (21 hpf), or differentiation (48 hpf), and all were fixed at 3 dpf, as schematized in **Figure 6A**. Embryos were assessed for enteric neuron abundance and localization via wholemount IHC, where Elavl3/4^+^ cells were counted along the gut tract (same region as in **Figure 5A, box**). When heat shocked at 14 or 21 hpf, GFP-Hoxb5b^+^ embryos formed significantly fewer Elavl3/4^+^ cells, when compared to their GFP-Hoxb5b^-^ sibling heat shock controls (**Figure 6B-E, H**). After heat shock at 48 hpf, GFP-Hoxb5b^+^ embryos did not significantly vary in number of Elavl3/4^+^ cells (**Figure 6F-H**). The distribution of Elavl3/4^+^ cells was weighted more heavily toward the midgut, though cells could be detected along the entire length of the gut in GFP-Hoxb5b^+^ embryos. This genetically encoded elevation of Hoxb5b activity during early NCC developmental phases corroborated the abrogation in Elavl3/4^+^ cells resulting from *vp16-hoxb5b* mRNA injection. Overall, these data indicate that the ability of GFP-Hoxb5b to affect enteric neuron number is limited to early stages of NCC development, but not thereafter.

**Figure 6.**
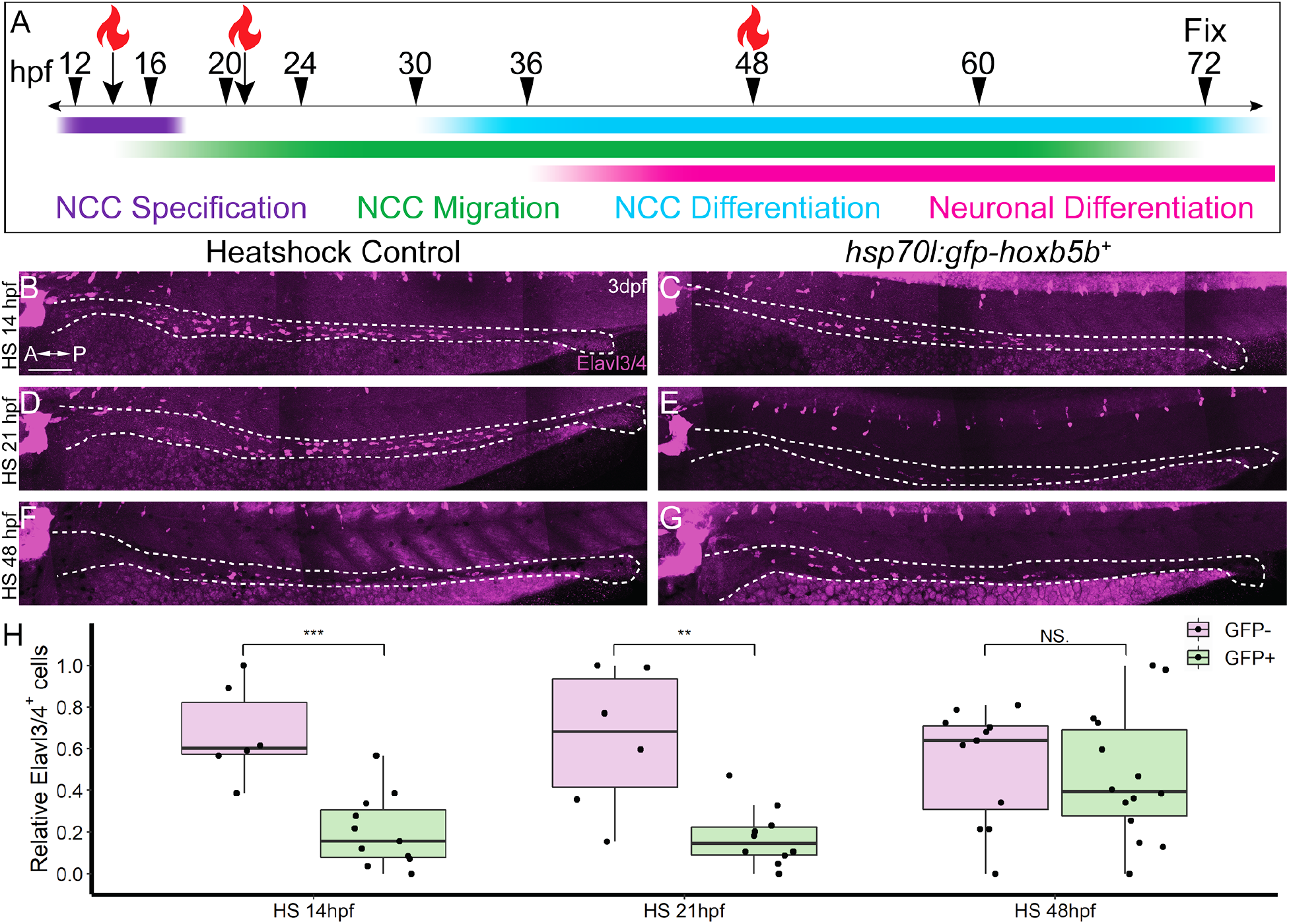
NCC sensitivity to increased Hoxb5b activity is restricted to earlier stages of development. **(A)** Schematized model of when heat shocks occurred relative to standard stages of NCC development in zebrafish. **(B-H)** Whole mount immunolabeled *hsp70l:GFP-hoxb5b*^*+*^ embryos and their GFP-sibling controls using an antibody against Elavl3/4. Embryos heat shocked either at 14 hpf or 21 hpf both exhibited fewer differentiating enteric neurons than controls. Numbers of enteric neurons are quantified in (H, HS14: *p* = 0.000232; HS21: *p* = 0.001684; HS48: *p* = 0.5813). Scale Bar B: 100 μm

### Excess Hoxb5b leads to stalled enteric nervous system development

We had thus far discovered that Hoxb5b was sufficient to strongly increase NCCs at 30 and 50 hpf, but suppressed the number of enteric neural progenitor cells by 3 dpf. The loss in cells could easily be explained by an acute wave of cell death during enteric neural progenitor migration. We tested this hypothesis with whole mount IHC probing for activated Caspase-3, a marker for apoptotic cells (Sorrells et al., 2013), as well as Phox2b to label enteric neural progenitors. In addition, we also conducted these experiments in the Tg*(−4*.*9sox10:EGFP*) embryos (hereafter referred to as *sox10:GFP*) (Carney et al., 2006), which marks migratory NCCs with cytoplasmic GFP, and relying on residual GFP signal post fixation to label the recently *sox10*^+^ enteric neural progenitor cells, as we have previously (Howard et al., 2021). Notably, Caspase-3^+^ cells were rare in all controls tissues examined from 33 to 66 hpf (**Figure 7 – Supplement 1A**,**C**,**E**,**G**). While small patches of apoptotic cells can be found proximal to the POD at 33 hpf and 55 hpf (**Figure 7 – Supplement 1B**,**D**), there was not a detectable onset of cell death between 55 and 66 hpf along the entire vagal and gut region (**Figure 7 – Supplement 1F**,**H**) to support the loss of enteric neural progenitors through apoptosis following Hoxb5b overexpression.

We next examined the progenitor state of enteric cells at 63 hpf, a window of development prior to the 3 dpf cut off, but after the 50 hpf NCC expansion noted previously. To this end, we asked if *sox10:GFP*^+^ and/or Phox2b^+^ cell numbers were reduced along the gut at 63 hpf. Intriguingly, *vp16-hoxb5b* did not lead to a significant change in the total number of GFP^+^ cells along the gut tube at 63 hpf (**Figure 7A-C**). Similarly, there was no change in the number of Phox2b^+^ cells, compared with control embryos (**Figure 7C**). This was in contrast to our earlier observation that enteric Phox2B^+^ cells were increased at 50 hpf (**Figure 5**). Furthermore, the fraction of *sox10:GFP*^+^ cells expressing Phox2b was roundly unchanged (**Figure 7D**). Therefore, we found that by 63 hpf the enteric progenitors exhibited enteric differentiation capacity, despite their decreased abundance in the presence of excess Hoxb5b.

**Figure 7.**
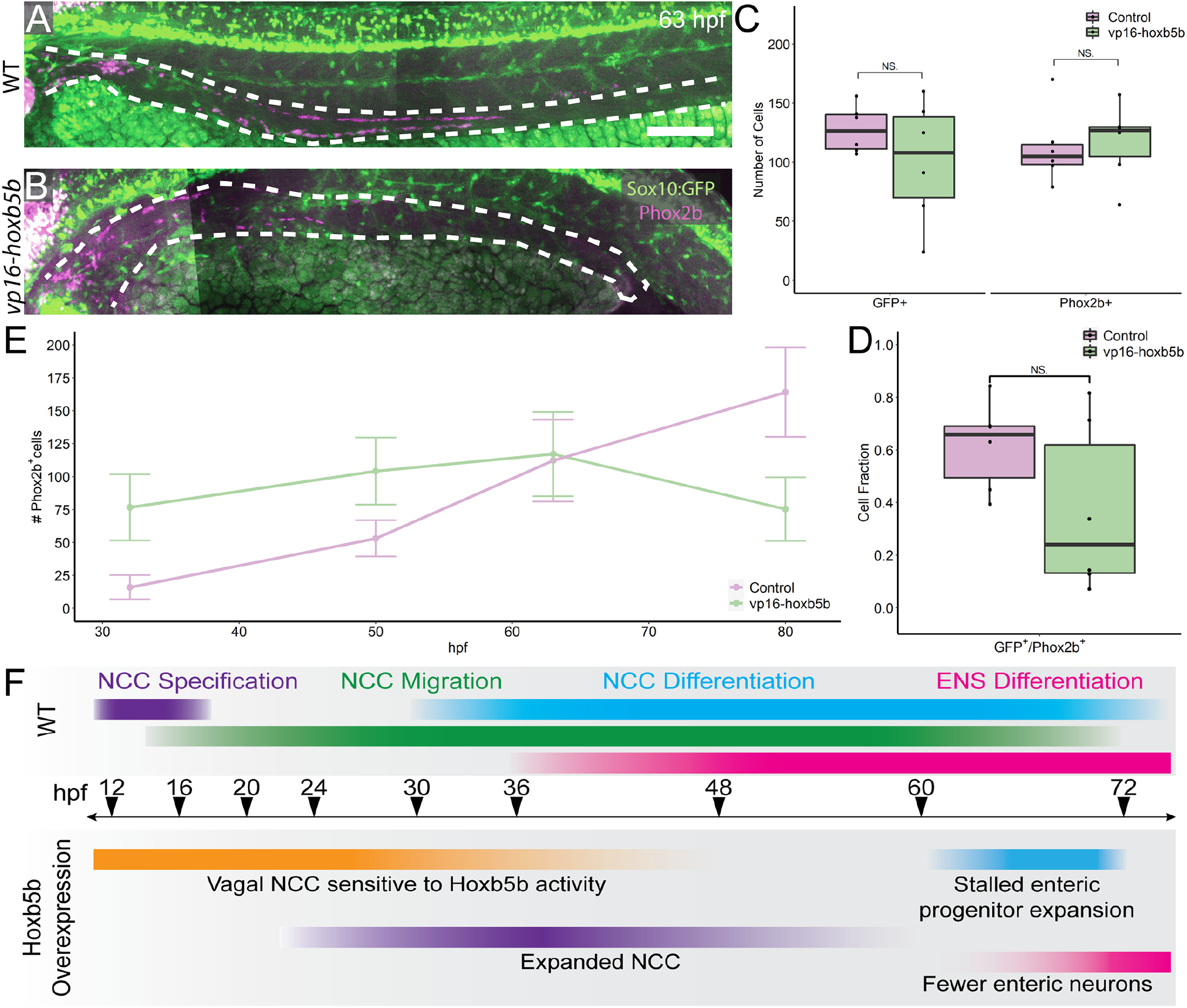
Elevated Hoxb5b abrogates the expansion capacity of enteric NCC-derived neuronal lineage. **(A-B)** By 63 hpf, *sox10:GFP*^*+*^ embryos immunolabeled for Phox2b show cells which have partially migrated along the gut in both control and *vp16-hoxb5b* injected animals. Gut tract outlined with white dashes. **(C)** No changes were found in the number of GFP^+^ NCC lineage or Phox2b^+^ cells along the gut (Enteric GFP^+^: *p* = 0.2619; Enteric Phox2b^+^: *p* = 0.7896). **(D)** Assessment of fraction of GFP^+^ cells which are also positive for Phox2b^+^ restricted to the gut axis, as a measure of NCC which have initiated their differentiation programs shows the fraction of So×10^+^ lineage cells which have turned on Phox2b expression is unchanged following elevated Hoxb5b (*p =* 0.1253). **(E)** Summary of the number of Phox2b^+^ cells for either WT control embryos or embryos injected with 15 pg *vp16-hoxb5b* mRNA as a function of age. The trend shows while the number of Phox2b^+^ cells initially is greater than WT, eventually the WT Phox2b^+^ numbers continue to increase with the Phox2b^+^ population in Hoxb5b elevated embryos remains flat. Error bars reflect ± one standard deviation. **(F)** Graphical Model of the role of Hoxb5b in NCC development, such that early Hoxb5b expression grossly expands NCC numbers, while later in development Hoxb5b suppresses NCC enteric expansion. Scale bar A: 100 μm

That we observed increased Phox2b^+^ cells at 32 and 50 hpf, yet saw a dampening of their numbers by 63 hpf, this suggested that the kinetics of enteric progenitor expansion may have been adversely affected following elevated Hoxb5b. Plotting the total number of Phox2b^+^ cells counted from each developmental stage assayed throughout this study (**Figure 7E**) revealed a steady increase in control Phox2b^+^ cells with developmental age, whereas Hoxb5b overexpressing animals present a stalled curve at 63 hpf. As we previously have shown that the enteric cells are not cleared by apoptosis during the 63-80 hpf transition, these results indicated that Hoxb5b activity modulates enterically-fated NCC capacity to expand as a population along the gut. The Hoxb5b-dependent precocious expansion of NCC leads to a fixed number available progenitors which are then unable to expand in sufficient numbers to lead to proper ENS formation. These findings position Hoxb5b as a fine-scale regulator of enteric NCC number.

## Discussion

We discovered that throughout the course of NCC specification and migratory phases, *hoxb5b* is expressed within subsets of vagal NCCs along the post-otic/posterior zebrafish embryo. Enhancement of Hoxb5b activity was sufficient to dramatically expand NCC localization patterns, as well as their number, along the embryo. The expansion of NCCs was also accompanied by domain expansions in vagal NCC marker genes *phox2bb* and *foxd3*. Temporally-restricted pulses of ectopic Hoxb5b during early vagal NCC developmental phases was sufficient to swiftly expand vagal NCC populations, which also persisted well into the second day in development. While many vagal NCC derivatives were unaltered by 3 dpf, NCC derivatives along the enteric neural trajectory were dramatically impacted. While we observed an increase in enteric neural progenitors along the developing gut tube at 50 hpf, they failed to expand and colonize the gut efficiently, resulting in a marked decrease in the number of enteric neurons. The decrease in the number of enteric neural progenitors following Hoxb5b induction appears to be due to a Hoxb5b-dependent modulation of the colonization capacity of enteric neural progenitors. Cumulatively these data position Hoxb5b as a potent regulator of NCC patterning and number during early embryonic development (**Figure 7F**).

The potential involvement of Hoxb5b in zebrafish NCC development has been suggested by previous expression analyses, with discrete expression along the dorsolateral neural tube and in the post-otic domain, posterior to rhombomere 8 (Barsh et al., 2017; Jarinova et al., 2008; Kudoh et al., 2001; Waxman et al., 2008). Additional expression domains are found within the lateral plate mesoderm and foregut by 24 hpf (Dalgin and Prince, 2021). As identified in a single cell atlas of posterior zebrafish NCC lineages at 48-50 and 68-70 hpf, *hoxb5b* was among the most pervasively expressed transcripts encoding for a Hox transcription factor in both the NCC and in neural fated lineages (Howard et al., 2021). Extending this prior work, our HCRs show for the first time at high resolution the persistent expression of *hoxb5b* in zebrafish posterior (post-otic) NCC through the early course of their development.

In our zebrafish model, despite the regional potential exhibited by NCCs (Rocha et al., 2020), elevated Hoxb5b uniformly expanded both cranial, vagal, and trunk NCC progenitors, though more robustly among the posterior NCC populations. Whether the Hoxb5b-dependent increase in NCCs is driven by increased NCC specification from the neural tube or through upregulation of NCC proliferation remains unresolved and was beyond the current scope of this study. Regardless of the underlying mechanism, these findings clearly indicate that Hoxb5b participates in NCC development from an early position in the NCC gene regulatory network (Martik and Bronner, 2017; Simoes-Costa and Bronner, 2016). Further, the rapid sensitivity of vagal NCC to temporally restricted pulses of elevated Hoxb5b suggests competency of these cells to abruptly respond to Hox activity early in their development. Indeed, early NCCs appear sensitive not only to Hox mediated activity but also the amount of Hoxb5b present. In our injection experiments, POD NCCs increased coordinately with amount of *vp16-hoxb5b* mRNA delivered. As such, the number of cells fated in select NCC lineages, such as the enteric NCCs, appear to be influenced not only by the activation of Hoxb5b-dependent activity, but also in part through the levels of Hoxb5b expression. From data derived across multiple animal models, Hoxb5b and its orthologues are known to be under control of several classical morphogenic signals, including WNTs (Lengerke et al., 2008), NOTCH (Hortopan and Baraban, 2011), and Retinoic Acid (Waxman et al., 2008), which enables fine spatiotemporal tuning of the levels of Hox expression. Summarily, our data thus illuminate a model in which Hoxb5b serves a potent regulator of posterior NCC identity and cell number, dependent on additional unspecified cofactors as well as its expression level.

Our findings regarding Hoxb5b in zebrafish NCC is complementary to and extends the developmental understanding of mammalian Hoxb5. For example, dominant negative suppression of Hoxb5 activity in NCC lineages led to a depletion of several NCC-derived cell populations including DRGs, pigment cells, as well as enteric neural progenitors (Kam et al., 2014b; Lui et al., 2008). Complementing these prior loss of function studies, our gain of function data indicates that elevated Hoxb5b activity is sufficient to induce expansion of vagal NCC progenitors—that paradoxically also leads to severe ENS hypoganglionosis. While we did not observe corresponding changes in DRG and pigment populations in our experiments; early enteric progenitors were dramatically increased in response to Hoxb5b activity along the gut, prior to the onset of neurogenesis, yet failed to properly execute enteric neuronal differentiation. The insufficiency of ectopic Hoxb5b to expand the DRG and pigment lineages suggests a separate regulatory mechanism for Hoxb5b gain of function activity in enteric NCC. Indeed, the specificity of the effect elevated Hoxb5b on NCC populations; despite global elevated Hoxb5b activity in both the heat shock and mRNA injection assays, only ENS progenitor pool was roundly perturbed, indicating a particular receptivity of this population to Hoxb5b transcriptional regulation. One possible frame work to explain this phenomena is the presence of additional co-factors, such as other Hox transcription factors and their regulators, within the ENS linage which cooperate with Hoxb5b to facilitate this specificity. Intriguingly, overexpression of murine Hoxa4, another proximally expressed Hox transcription factor, is also endemic to the gut nervous network and is known to result in megacolon when over expressed, a phenotype indicative of enteric neural crest disruption (Tennyson et al., 1993; Wolgemuth et al., 1989). Reflecting on the emerging numerous descriptions of combinatorial “Hox Codes” which define NCC identity (Howard et al., 2021; Parker et al., 2019; Soldatov et al., 2019), the prospect of a shared sensitivity in enteric NCCs to Hoxb5b and Hoxa4 becomes a tantalizing speculation. When taken together with mammalian suppression of activity studies referenced above, our gain of function results suggest that the vertebrate embryo is exquisitely sensitive to perturbations in Hoxb5 activity, where either elevations or reductions in Hoxb5 lead to severe ENS defects.

Somewhat counterintuitively, the increase in POD NCCs following increased Hoxb5b activity did not correspondingly manifest a pan increase in vagal-derived NCC lineages. While many vagal-derived lineages exhibited no discernable phenotypic change, we actually observed a dramatic decrease in the number of enteric neurons in animals with elevated Hoxb5b activity. The shift in abundance of enterically fated cells was not the result of NCC-specific cell death or abrogated differentiation potential; indeed there was no change in fractions of enteric NCCs which had initiated differentiation programs following elevated Hoxb5b. Rather this tissue specific cell decrease appears to be the caused by a late-onset suppression of enteric neuroblasts expansion. These results suggest that Hoxb5b not only participates in regulating vagal NCC number over the influence of fate acquisition in this subset of NCC lineages. We find it tempting to speculate that perhaps the failure to perturb fate acquisition is at least in part mediated through functional redundancy of other Hox genes (Jarinova et al., 2008) as well as other factors, such as *foxd3*, which was grossly expanded following Hoxb5b ectopic activation. *Foxd3* has previously been identified as a known transcriptional target (Kam et al., 2014a) and further modulates fate acquisition of several NCC lineages (Lukoseviciute et al., 2018). As the downstream mechanism remains to be elucidated, our findings here demonstrate for the first time that Hoxb5b is sufficient to grossly abrogate the expansion capacity of a very specific niche of NCCs, which suggests an exciting model for the role of additional Hox factors within the context of posterior NCCs. Collectively, our findings are in support of a model in which Hoxb5b plays an important role in NCC development, demonstrating the capacity to both expand vagal NCC localization and numbers. While additional questions still remain, these findings greatly inform our understanding of the role of posterior Hox genes in NCC development.

## Methods

### Zebrafish Husbandry and transgenic lines

Synchronously staged embryos for each experiment were collected via controlled breeding of adult zebrafish. After collection, embryos were maintained in standard E3 media at 28°C until 24 hours post fertilization (hpf), then transferred to 0.003% 1-phenyl 2-thiourea (PTU)/E3 solution (Karlsson et al., 2001), with the exception of larvae used to assay pigmentation, which were cultured in E3 media only. Transgenic embryos for the Tg(*-4*.*9sox10:EGFP*)^ba2Tg^ (Carney et al., 2006) and Tg(*-7*.*2sox10:mRFP*)^vu234^ (Kucenas et al., 2008) were generally sorted between 17-28 hpf for fluoresces while Tg(*hsp70l:EGFP-hoxb5b;acry:dsRed*)^ci1014^ and Tg(*-8*.*3phox2bb*:*kaede*) (Harrison et al., 2014) embryos were sorted for transgenic expression between 60-78 hpf. Tissue was collected from embryos out of their chorions at the stage noted in each experiment as described in (Ibarra-García-Padilla et al., 2021). All work was performed under protocols approved by, and in accordance with, the Rice University Institutional Animal Care and Use Committee (IACUC).

### Generation of the *Tg(hsp70l:EGFP-Hoxb5b)* transgenic line

To generate EGFP-Hoxb5b, *egfp* was fused to the 5’-end of zebrafish *hoxb5b* with PCR. A sequence encoding a 7 amino acid linker was incorporated and the *hoxb5b* ATG was deleted to prevent alternative transcriptional initiation of *hoxb5b* downstream. To generate the *hsp70l:EGFP-Hoxb5b* transgene, standard Gateway methods were used (Kwan et al., 2007). The transgene includes a *EGFP-Hoxb5b* middle-entry vector and the reported *p5E-hsp70l* 5’-Entry and *p3E-polyA* 3’-entry vectors (Kwan et al., 2007), which were incorporated into the *pDestTol2-acry:dsRed* vector (Mandal et al., 2013). Sanger sequencing was used to confirm the proper orientation of the constructs within the destination vector and the sequence of *EGFP-hoxb5b*. Transgenic embryos were created by co-injecting wild-type embryos at the one-cell stage with 25 pg *hsp70l:EGFP-hoxb5b* vector and 25 pg of *Tol2* mRNA (Kawakami, 2004; Kawakami et al., 2004). Embryos were raised to adulthood and screened for the present of dsRed in the lens at ∼3 days and the ability to induce robust EGFP expression following heat-shock (**Figure 4 – Supplement 1**). Multiple founders for the Tg(*hsp70l:EGFP-hoxb5b;acry:dsRed*)^ci1014^ line were identified so the line that induced the most robust expression following heat-shock was retained. While some ectopic notochord expression was observed in non-heat shocked embryos they did not exhibit overt phenotypes and developed normally. For heat shock experiments, through routine outcrossing of transgenic animals to the wild type embryos of the AB/TL backgrounds, GFP-hoxb5b^-/-^ siblings are produced with each subsequent breeding, which were heat shocked and processed in parallel.

### Preparation & Injections of *hoxb5b* mRNA

Capped *vp16-hoxb5b* mRNA was prepared off a Not1 linearized pCS2+ plasmid containing the *vp16-hoxb5b* coding sequence using the Sp6 mMessage Kit (Ambion), as first reported in (Waxman et al., 2008). The *vp16-hoxb5b* construct encodes for a hyperactive form of Hoxb5b and allowed for lower doses of mRNA to be delivered (Waxman and Yelon, 2009). Embryos were injected prior to the four-cell stage with either 15 pg or 30 pg of mRNA, which were cultured in parallel with uninjected siblings used for controls. Dead and grossly malformed embryos were removed from analysis.

### In Situ Hybridization

In situ hybridizations were performed similarly to the protocol of Jowett and Lettice, 1994, which should be referenced for specific details. Briefly, antisense digoxigenin-labeled riboprobes were generated from previously characterized plasmids containing sequences for *crestin* (Luo et al., 2001), *foxd3* (Hochgreb-Hagele and Bronner, 2013; Odenthal and Nüsslein-Volhard, 1998), *phox2bb* (Uribe and Bronner, 2015), and *hoxb5b* (Waxman et al., 2008). As per the protocol, whole mount embryos stored in methanol were rehydrated in PBST, permeabilized with Proteinase K digestion (10 μg/ml), and refixed in 4% PFA. Embryos were incubated in probes overnight (∼16 hours) at 65°C and washed sequentially in graded SSCT buffers. Riboprobes solutions were recovered and stored at -20°C for reuse, with multiple uses leading to minimal loss of signal. Probed embryos were blocked for 1-2 hours at ambient temperature in 5% Goat sera in PBST before detection overnight (∼16 hours) at 4°C using an anti-Digoxigenin-Fab fragments conjugated to Alkaline Phosphatase enzymes (1:1000 dilution, Roche) in 5% Goat Sera in PBST. Finally, riboprobes were visualized with NBT/BCIP solution (3.5 μL each of NBT, BCIP stock solutions, Roche). Probes were validated prior to use on wildtype embryos to calibrate staining duration, with patterns compared to those curated on ZFIN (Howe et al., 2013; Ruzicka et al., 2019).

### Whole mount Immunohistochemistry, HCR, and WICHCR

Immunohistochemistry (IHC), Hybridization Chain Reaction (HCR), and Whole mount Immuno-Coupled Hybridization Chain Reaction (WICHCR) protocols all were conducted according to the methods published in Ibarra-García-Padilla et al., 2021. All IHC assays conducted in blocking 5% Goat Sera in 1X PBST. Primary antibodies against the following proteins were used as follows: Phox2b (1:200, Santa Cruz, B-11), Kaede (1:500, MBL International, PM102M), Elavl3/4 (Same as HuC/D, 1:500, Invitrogen Molecular Probes, A21271), Activated Caspase-3 (1:200, BD Biosciences, 559565). Incubation in primary antibody solutions were conducted overnight at 4°C, except for assays with Phox2b or Caspase-3 antibodies which were allowed to incubate for two days at 4°C which provided optimal labeling. Corresponding secondary antibodies conjugated to spectrally distinct fluorophores were all used at 1:500 dilution, selected from the following depending on the experimental condition: Alexa Fluor 488 goat anti-rabbit IgG (ThermoFischer, A11008), Alexa Fluor 568 goat anti-rabbit IgG (ThermoFischer, A11011), Alexa Fluor 488 goat anti-mouse IgG1 (ThermoFischer, A21121), Alexa Fluor 594 goat anti-mouse IgG1 (ThermoFischer, A21125), Alexa Fluor 647 goat anti-mouse IgG1 (ThermoFischer, A21240), and Alexa Fluor 647 goat anti-mouse IgG2b (ThermoFischer, A21242). In the HCR & WICHCR assays, commercially designed probes were secured from Molecular Instruments as follows: *crestin* (B3, AF195881.1), *hoxb5b* (B2, BC078285.1), *phox2bb* (B1, NM_001014818.1). Corresponding amplifiers were purchased from Molecular instruments and were used in experiments to include spectrally distinct fluorophores suitable for multiplexed imaging.

### Heat shock induction of the *hsp70l:GFP-hoxb5b* transgene

Adult zebrafish maintained as an outcross and positive for dsRed expression as larvae in the lens (see above method on description of the line) were bred to produce synchronously staged embryos. At the stage designated to begin the heat shock, embryos were rapidly transferred to 37°C E3 and maintained at that temperature. After a 1-hour incubation, embryos were rapidly returned to 28°C. In the 1-3 hours after heat shock GFP-Hoxb5b^+^ embryos were sorted from GFP^-^ siblings and cultured in parallel until the designated stage for tissue collection.

### Imaging, Quantification, and Image Visualization

All embryos prior to imaging were cleared through graded washes of PBST/Glycerol to reach a final Glycerol content of 75%. Fluorescent Z-stacked images of IHC processed embryos were captured using an Olympus FV3000 point scanning confocal microscope supported by Fluoview Acquisition Software (version FV31S-SW). Images were stitched in FIJI (Image J version 1.53e) using the Grid/Collection applet as part of the Stitching plugin (Rueden et al., 2017; Schindelin et al., 2012; Schneider et al., 2012). Digital image files were converted with ImarisFileConverter Software (Bitplane) to three dimensional rendered images compatible with IMARIS (V9.4, V9.7, Bitplane). Cells counts were conducted on volume images following an arithmetic background subtraction in IMARIS to ensure accurate counts, particularly when determining coincidence of labels. ISH processed embryos were imaged on a Nikon Eclipse Ni microscope equipped with a motorized stage. Z-stack images were acquired and extended depth of focus images were generated in the Nikon NIS-Elements BR software (v5.02.00). Areas of expression were measured in FIJI to include dark pixels in the post-otic or hindbrain domains. Quantifications were curated and analyzed in the Rstudio programming environment (v1.1.463). Images of pigmented embryos were captured similarly on the Nikon microscope with lateral illumination to distinguish iridophores, similar to (Petratou et al., 2021).

### In vivo Confocal Microscopy

Embryos were sorted for RFP expression, anesthetized with 0.4% tricaine, and embedded in 1% low melting agarose in a 28.5 °C chamber with a coverslip glass bottom. Care was taken to ensure embryo angled appropriately and proximal to the glass. Z-stack images were acquired approximately every half hour concurrently on the same Olympus FV3000 point scanning confocal microscope as above. Maximum intensity projections images were generated and exported from FIJI.

### Statistical Analysis

An α of 0.05 was used as a cut off for all statistical tests. Normalcy of datasets was assessed by visual inspection of a density plot, a qqplot against a linear theoretical distribution, and Shapiro-Wilk test for Normalcy. Further, variance between each dataset was examined either with a Bartlett test for data which adhered to normalcy or with a Levene’s test for non-normal data. Based on the normalcy and scedasticity conditions of the data, the appropriate statistical test was selected, as summarized in Table 1. All statistical analyses were carried out in the Rstudio (v1.1.463) programming environment, with key dependencies on the lawstat (v3.4) and stats (v3.6.3) packages. Plots were generated in Rstudio supported by the ggplot2 (v3.3.2) and ggsignif (v0.6.0) packages.

## Figure Legends

**Figure 3–Supplement 1.**
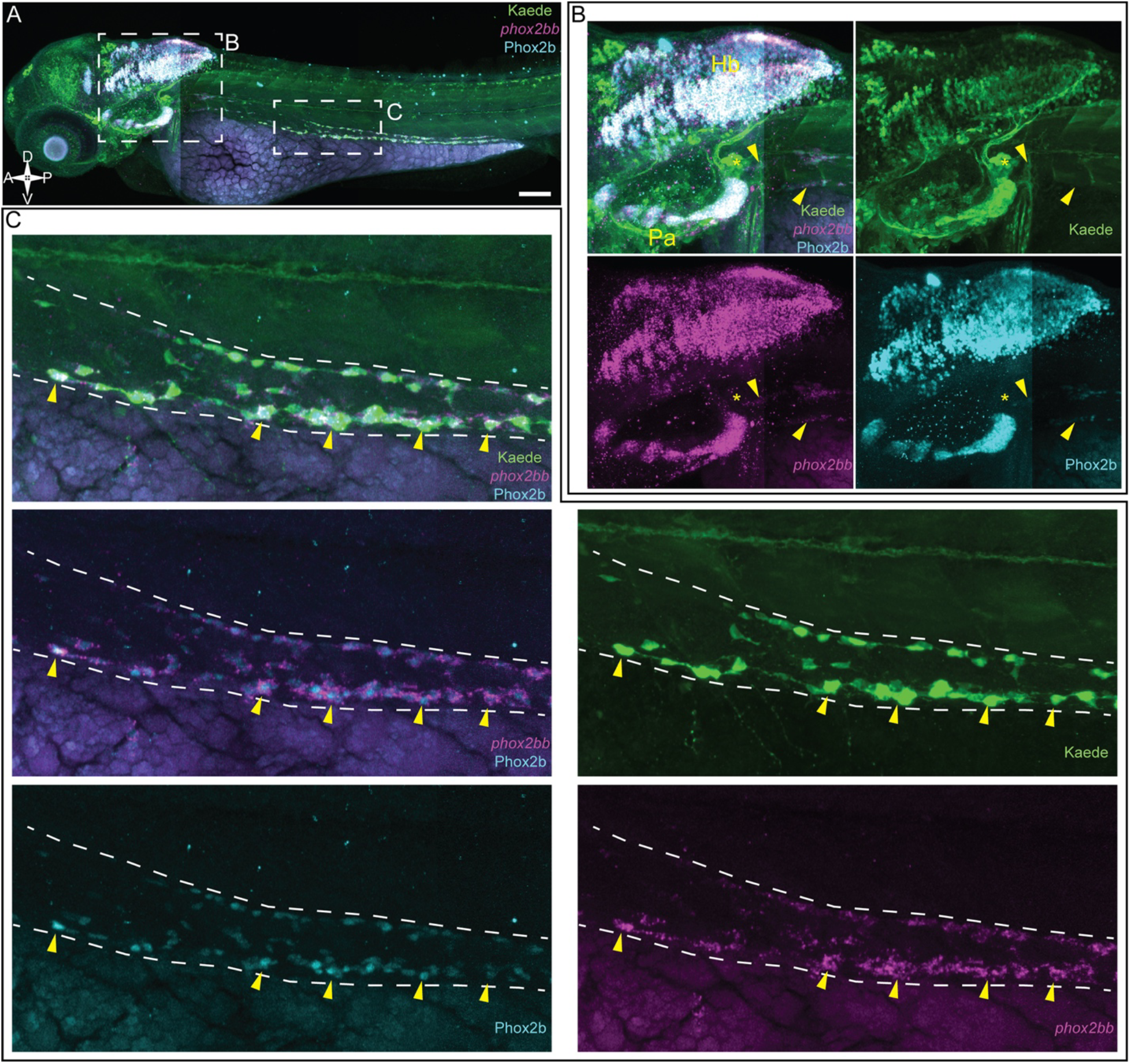
Phox2b antibody labeling is synonymous with mRNA & transgenic labeling methods. **(A)** 3 dpf *-8*.*3phox2bb*:*kaede* larva co-labeled via WICHCR protocol (Ibarra-García-Padilla et al., 2021) showing transgene labeled cells (green), *phox2bb* mRNA (magenta), and Phox2b protein (cyan). **(B)** Inset images in the hindbrain exhibit robust coincidence of all three probes along the Hindbrain (Hb), Pharyngeal (Pa), and PODs (arrowheads). Kaede^+^ cells which are negative for both mRNA and protein are detected at the dorsal most aspect of the motor neuron complex (star) as well as in anterior regions of the presumptive CNS. **(C)** Images highlighting the midgut axis reveals a perfect coincidence of all three enteric labels with very little background, demonstrating the efficacy of the Phox2b antibody as an enteric neural crest label. While nearly every cell is coincident, selected cells are annotated to aid in comparison (arrowheads). Scale bar A: 100 μm

**Figure 4-Supplement 1.**
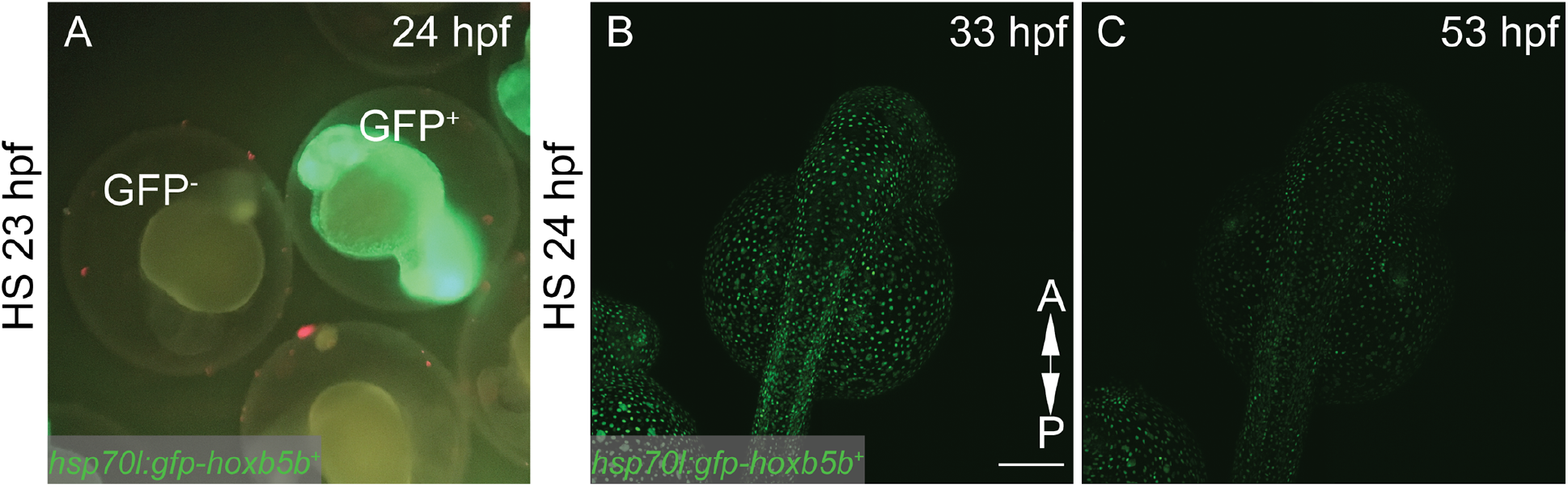
Characterization of novel *hsp70l:GFP-hoxb5b* transgenic zebrafish line. **(A)** GFP-Hoxb5b^+^ overexpressing embryos are easily distinguishable from GFP^-^ siblings one hour after heat shock. GFP^-^ & GFP^+^ embryos are cultured and assayed under identical conditions to control for variations induced by growth at an elevated temperature. **(B-C)** GFP-hoxb5b construct is robustly localized to cell nuclei following heat shock and persists at least a day after initial induction (C). Scale bar B: 100μM

**Figure 5-Supplement 1.**
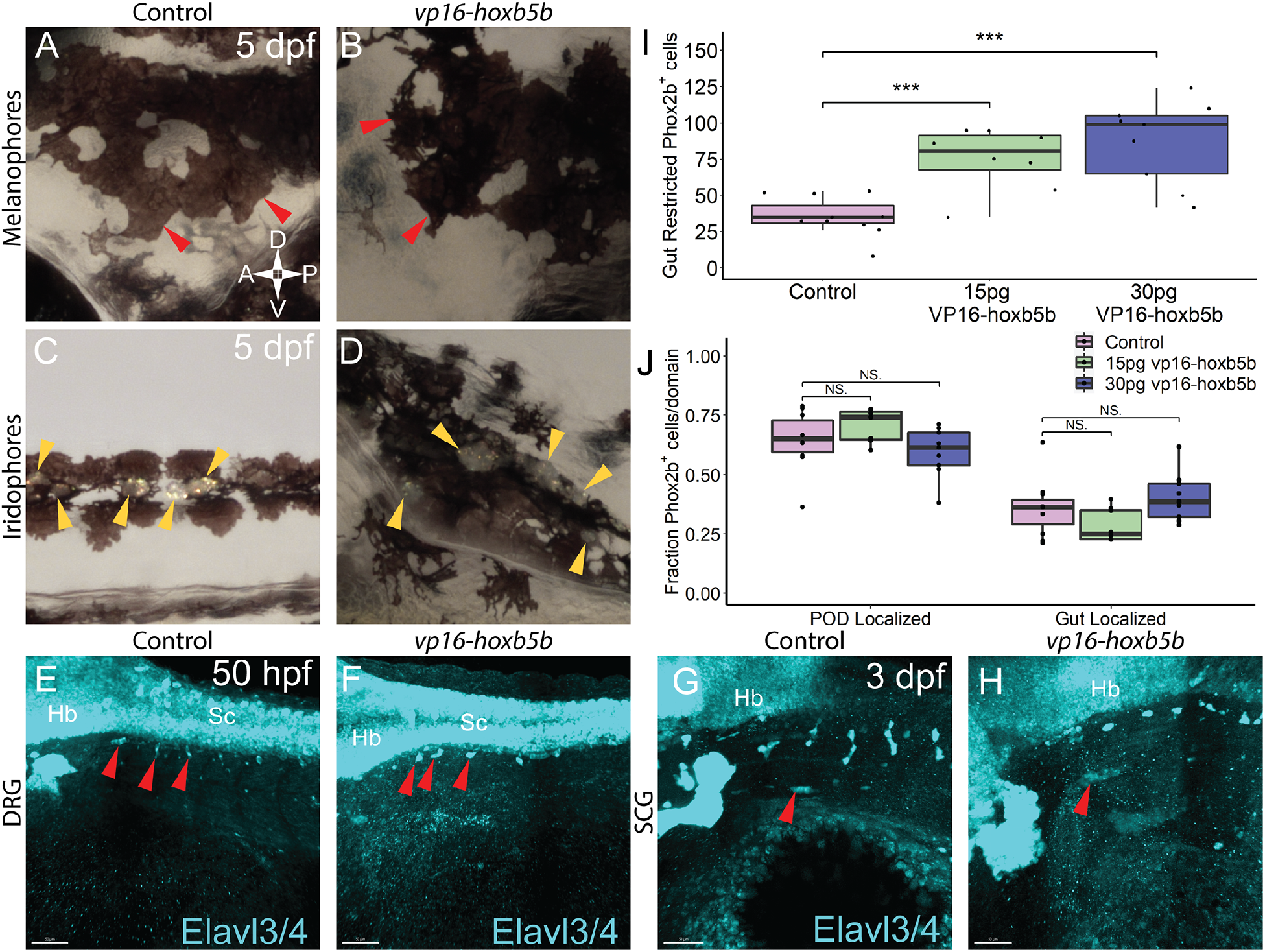
Vagal NCC derivatives following Hoxb5b overexpression are largely unaffected. **(A-D)** Investigation of distinct NCC derived pigment lineages, melanophores (red arrowheads) and iridophores (yellow arrowheads), in 5 dpf embryos overexpressing Hoxb5b compared to controls. **(E-F)** Both control embryos and Hoxb5b overexpressing embryos by 50 hpf produced phenotypically wild type anterior dorsal root ganglia (DRG), as shown in whole mount immunolabeling with an antibody against Elavl3/4. **(G-H)** Position and development of the Superior Cervical Ganglion (SCG) occurs by the third day, as shown by immune labeling for Elavl3/4 (red arrowhead). **(I)** Quantifications from data reflect in Figure 5, showing the number of Phox2b^+^ cells localized along the gut length, excluding the POD increases in response to *vp16-hoxb5b* mRNA induction (I, 15 pg: *p* = 9.92 × 10^−5^; 30 pg: *p* = 0.000404). **(J)** The distribution of Phox2b^+^ cells between the POD and the length of the gut was unchanged following elevated Hoxb5b activity (J, 15 pg in POD: *p* = 0.2515; 30 pg in POD: *p* = 0.3673; 15 pg in Gut: *p* = 0.2306; 30 pg in Gut: *p* = 0.3496). Scale Bars E,F,G,H: 50 μm

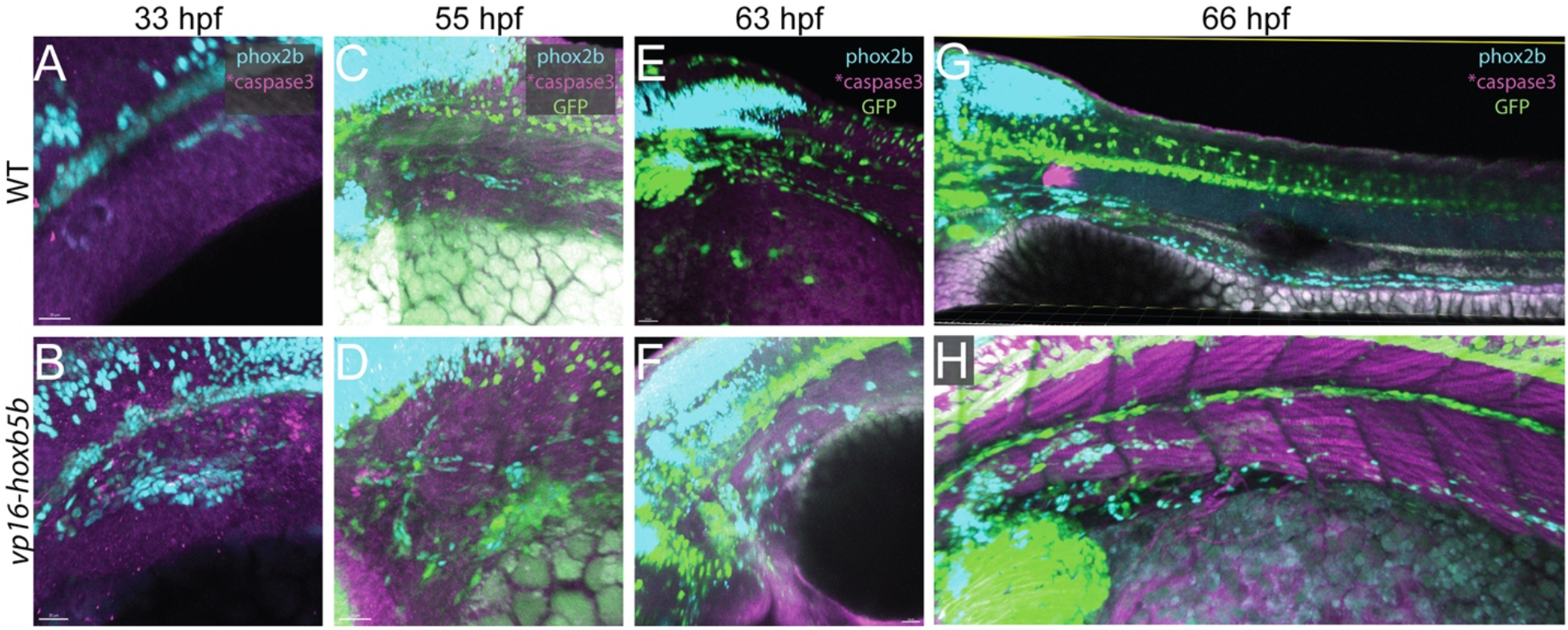

## Author Contributions

AGA-H and RAU designed the experiments and wrote the manuscript. RAU and AGA-H conceived the idea. AGA-H, ACN, JT, PR, EWS, CL, GK and RU performed the experiments. PR and JSW created the zebrafish transgenic line. AGA-H and RAU performed the data analyses. All the authors contributed to the article and approved the submitted version.

## Funding

Funding for this project was provided by Rice University, a Burroughs Wellcome Fund PDEP Award, Cancer Prevention & Research Institute of Texas (CPRIT) Recruitment of First-Time Tenure Track Faculty Members (CPRIT-RR170062) and the NSF CAREER Award (1942019) awarded to R.A.U as well as through the NIH NHLBI (HL137766, HL141186) awarded to J.S.W.

## Acknowledgements

We offer our sincerest gratitude to Dr. Dan Wagner and to the entire Uribe Lab at Rice University for their insights and support throughout this project. We thank Dr. Budi Utama and the Rice University Shared Equipment Authority on IMARIS image analysis suite, which was indispensable toward this project. We also thank Dr. Eric Bridenbaugh and Dr. Mariane Martinez at Olympus for their expert advice on confocal microscopy.

## Notes

### Competing Interest Statement

The authors have declared no competing interest.

